# Canonical translation factors eIF1A and eIF5B modulate the initiation step of repeat-associated non-AUG translation

**DOI:** 10.1101/2025.05.19.654993

**Authors:** Hayato Ito, Kodai Machida, Yuzo Fujino, Mayuka Hasumi, Soyoka Sakamoto, Yoshitaka Nagai, Hiroaki Imataka, Hideki Taguchi

## Abstract

Nucleotide repeat expansions, such as the GGGGCC repeats in *C9orf72*, associated with C9-ALS, are linked to neurodegenerative diseases. These repeat sequences undergo a non-canonical translation known as repeat-associated non-AUG (RAN) translation. Unlike canonical translation, RAN translation initiates from non-AUG codons and occurs in all reading frames. To identify potential regulators of RAN translation, we employed a bottom-up approach using a human factor-based reconstituted cell-free translation system to recapitulate RAN translation. This approach revealed that omission of either eIF1A or eIF5B enhanced the translation in all reading frames of *C9orf72*-mediated RAN translation (C9-RAN), suggesting that eIF1A and eIF5B act as repressors of RAN translation. eIF1A and eIF5B are known to contribute to the fidelity of translation initiation. In HEK293T cells, double knockdown of eIF1A and eIF5B further promoted C9-RAN compared to single knockdowns, indicating that these factors regulate C9-RAN through distinct initiation steps. Furthermore, under eIF1A knockdown conditions, the enhancement of RAN translation via the integrated stress response (ISR) was not observed in HEK293T cells, indicating that eIF1A is involved in the ISR-mediated non-AUG translation.

**GRAPHICAL ABSTRACT:** 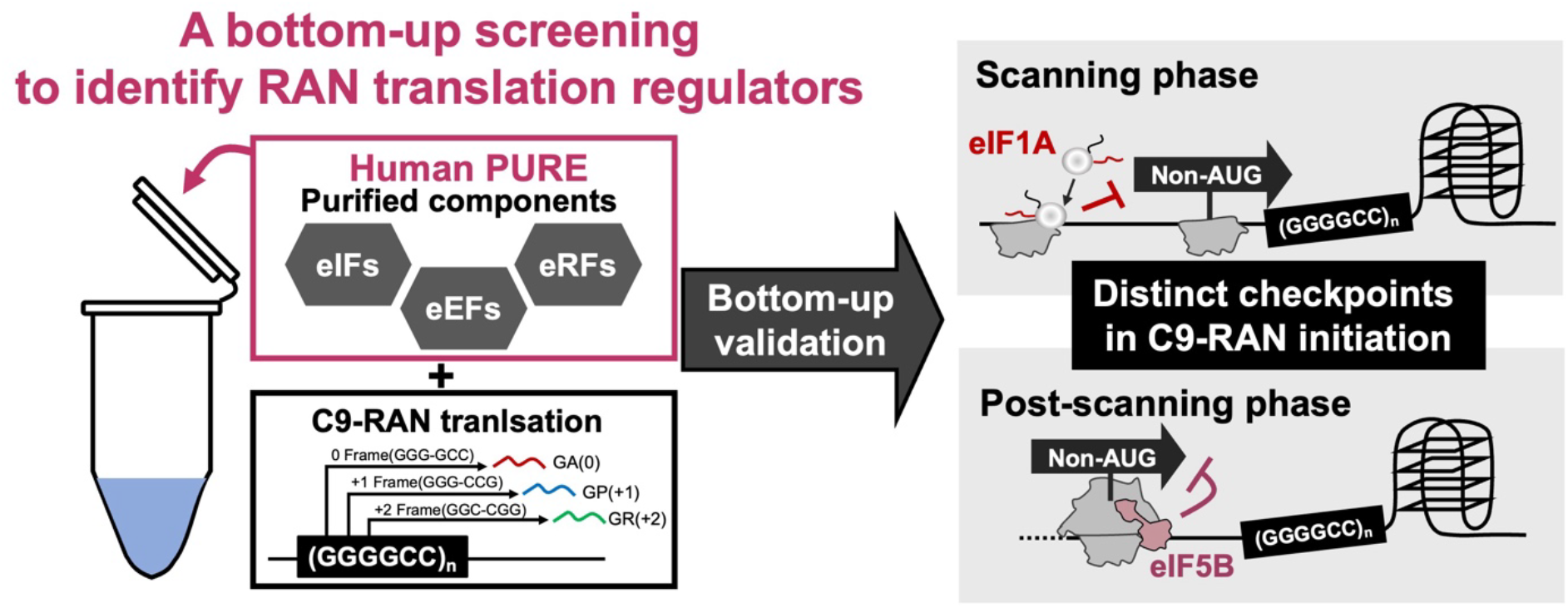

## INTRODUCTION

The human genome contains numerous repeat sequences. Expansion of certain repeats, such as GGGGCC in *C9orf72* and GGCCTG in *Nop56*, is associated with neurodegenerative diseases, including amyotrophic lateral sclerosis (ALS) and spinocerebellar ataxia (SCA) (1, 2). These repeat expansions disrupt splicing of their respective genes, resulting in transcripts that contain the expanded repeat (3–5). Recent studies have shown that repeat-associated non-AUG (RAN) translation, a process in which translation is initiated independently of the AUG start codons (6–9). A key feature of RAN translation is the expression on both sense and antisense transcripts, allowing translation in all possible reading frames (10–12). A well-studied example is RAN translation of the *C9orf72*-GGGGCC repeat (C9-RAN), which produces dipeptide repeats (DPRs) such as Gly-Ala (GA(0)), Gly-Pro (GP(+1)), and Gly-Arg (GR(+2)) from the sense strand (GGGGCC)_n_ repeat (7, 8, 10). These DPRs have been detected in patient-derived cells and various model organisms, where they exhibit toxicity (7, 8, 10, 12–16). Based on these findings, targeting the regulatory mechanisms of RAN translation may represent a novel therapeutic approach.

In eukaryotes, translation initiation occurs through a scanning mechanism (Fig. 1A) (17–19). Briefly, the 40S ribosomal subunit, along with initiation factors such as eIF1, eIF1A, eIF3, and the eIF4F complex (eIF4A, eIF4E, and eIF4G), binds to the 5’ cap of the mRNA and scans for an optimal start codon. eIF1 and eIF1A are crucial for this process by facilitating scanning and ensuring recognition of the optimal start codon (P_in_ state) (19). After scanning the start codon (post-scanning phase), translation factors, excluding eIF1A, dissociate from the scanning complex, and eIF5B, together with eIF1A, promotes the joining of the 60S subunit, forming the functional 80S ribosome and transition to elongation (20–22). The initiation mechanism of RAN translation shares several features with canonical translation pathways (Fig. 1A), particularly at the rate-limiting initiation stage (23–28). For example, C9-RAN, *Nop56* RAN translation (NOP56-RAN), and *Fmr1* CGG repeat RAN translation (CGG-RAN) initiate translation in a 5’ cap- and eIF4A-dependent manner, similar to canonical translation (23–28). These RAN translations initiate from near-cognate codons upstream of the repeat sequences through a conserved scanning mechanism. We previously demonstrated that C9-RAN can be recapitulated using a reconstituted 5’ cap-dependent human cell-free translation system containing minimal canonical translation factors (human PURE system) (28). These findings suggest that the core mechanisms of RAN translation largely overlap with those of canonical translation. In contrast, several distinct mechanisms of RAN translation have also been reported. Bicistronic reporter assays and single-molecule imaging studies suggest that C9-RAN can initiate translation via cap-independent mechanisms (29–32). Additionally, RAN translation is enhanced under conditions that activate integrated stress response (ISR), a cellular state that globally suppresses canonical translation through phosphorylation of eIF2α (24, 27, 30, 33). These observations indicate the presence of multiple translation initiation modes in RAN translation.

**Figure 1.**
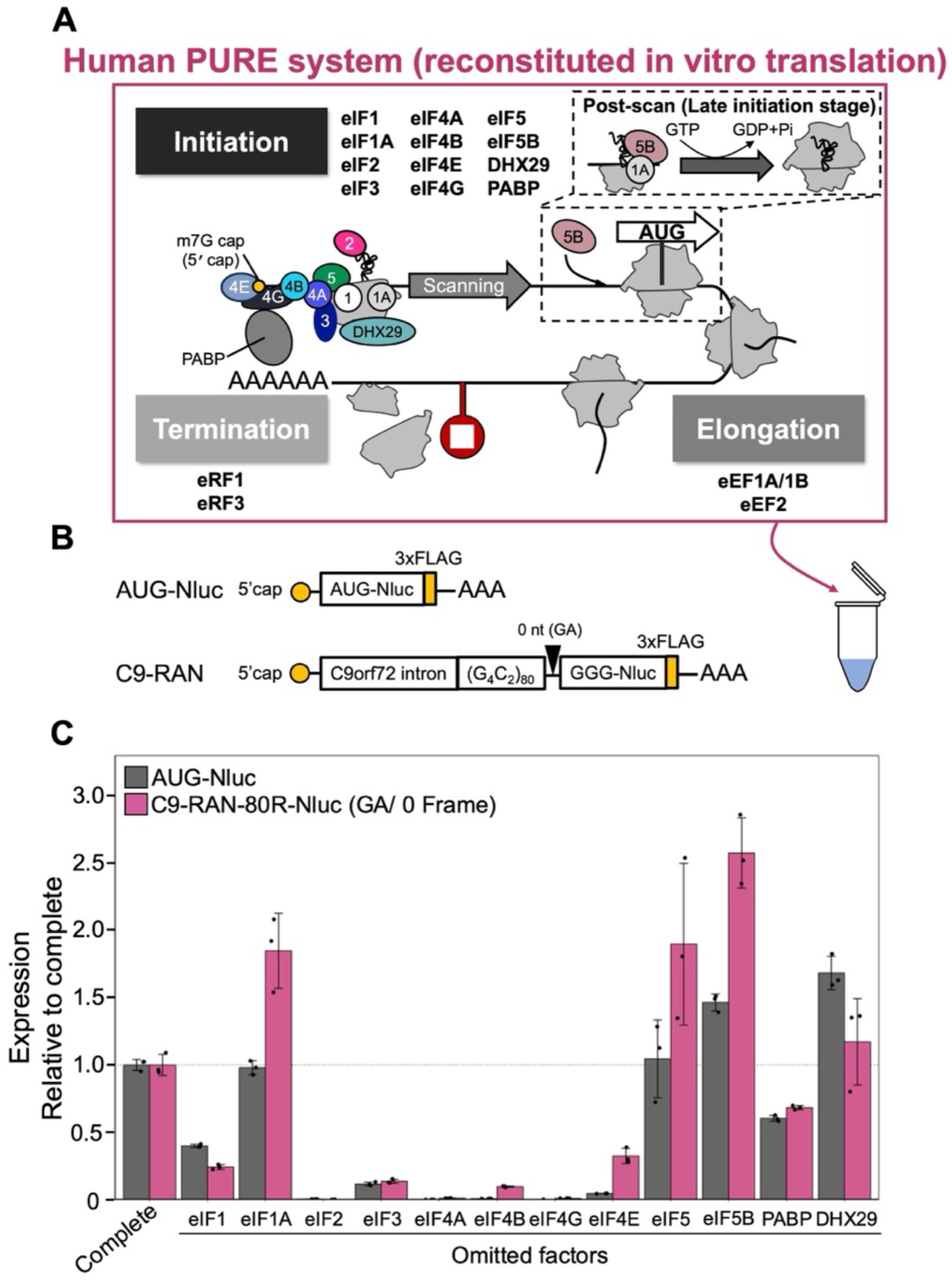
Human PURE system screening identifies eIF1A and eIF5B as repressors of C9-RAN. (**A**) Schematic of the human PURE system, a reconstituted translation system composed of purified translation factors (47, 48). (**B**) Schematic of the C9-RAN Nanoluciferase (Nluc) reporters, based on previously described constructs (24, 28). Nluc was used to quantify translation efficiency. (**C**) Relative expression of AUG-Nluc and C9-RAN reporters in the human PURE system upon systematic omission of the indicated translation factors. Values were normalized to the complete human PURE system (see Supplementary Fig. S1 for the experimental schematic). All graphs show mean ± SD from three independent experiments.

Recent studies have reported that canonical translation fidelity factors such as eIF1 and eIF5 act as regulators of CGG-RAN and C9-RAN (34, 35). In addition, alternative (non-canonical) initiation factors, including eIF2A, eIF2D, MCTS1/DENR, and DAP5, have been identified as regulators of both C9-RAN and CGG-RAN (31, 36–38). These factors are implicated in regulating translation initiation at non-AUG codons. However, our previous reconstitution of C9-RAN using the human PURE system showed that eIF2A and eIF2D did not promote RAN translation, suggesting that they either do not function independently or influence C9-RAN indirectly through secondary effects (28). Furthermore, several RNA-binding proteins, such as DDX3X, DHX36, and FUS, which interact with repeat RNA and modulate its structure, have also been identified as regulators of RAN translation (34, 39–42). Since non-AUG translation initiation has been reported to be facilitated by downstream RNA secondary structures (43, 44), we postulate that the fundamental mechanism of RAN translation involves altered 40S ribosome scanning dynamics caused by repeat RNAs, leading to misrecognition of the start codon.

The identification of RAN translation regulators has so far relied on comprehensive screening approaches, such as genome-wide CRISPR-Cas9 screens in cultured cells and *in vivo* screenings using *Drosophila* (34, 39, 45, 46). While these approaches are powerful for identifying novel RAN translation modulators, the species and cell types used, as well as secondary effects, may lead to the identification of incorrect factors. Translation-related factors are often essential genes, and their knockdown or knockout is likely to cause substantial secondary effects. For example, although DHX36 knockout was identified in a CRISPR screen as a candidate that suppresses RAN translation, further detailed analysis revealed that DHX36 actually promotes RAN translation (39–41).

To avoid these issues, here we employed a bottom-up screening approach using our previously developed human PURE to identify RAN translation regulators (28, 47, 48). The PURE-based screening allowed us to focus on the mechanisms without confounding secondary effects of essential factors, enabling the identification of specific regulatory components. Through this screening, we found that eIF1A and eIF5B function as RAN translation regulators. Both were shown to inhibit non-AUG translation initiation in RAN translation, as demonstrated by cell-free and cell-based assays. Moreover, under eIF1A knockdown conditions, the ISR-induced enhancement of RAN translation was abolished. These findings suggest that regulation of scanning and non-AUG translation initiation by canonical translation factors plays a key role in controlling C9-RAN initiation and in reprogramming C9-RAN under ISR conditions.

## MATERIAL AND METHODS

### Plasmids

For the in vitro translation assay, the reporter constructs T7-*C9orf72* intron1-(G_4_C_2_)_80_-nanoluciferase (Nluc)-3×FLAG, T7-ATG/CTG-Nluc-3×FLAG, and T7-NOP56-(GGCCTG)_71_-Nluc-3×HA were generated previously (27). For the in vivo translation assay, pcDNA™5/FRT-T7-*C9orf72* intron1-(G_4_C_2_)_66_-0/+1/+2-Nluc-3×FLAG was constructed in two steps, (i) pAG3-*C9orf72* intron1-(G_4_C_2_)_66_-3 frame tag (FT) (25) was digested by HindIII and NotI (NEB), and inserted into the pcDNA™5/FRT-T7 vector; (ii) the Nluc sequence was amplified by PCR and inserted into the pcDNA™5/FRT-T7-(G_4_C_2_)_66_ vector digested with NotI via Gibson Assembly. Different repeat lengths were generated randomly by PCR. The mutation of the CTG codon in the GA (0) frame to the ATG codon was carried out as described previously (28). Additionally, *C9orf72* intron1-AUG-Nluc-3×FLAG was constructed by inserting the *C9orf72* intron into an AUG-Nluc-3×FLAG reporter (28) via Gibson Assembly. For the in vivo overexpression assay, eIF1A and eIF5B coding sequences were amplified by PCR and inserted into a pAG vector containing a CAG promoter. Supplementary Tables S1 and S2 provide a detailed list of the plasmids and oligonucleotides used in this study.

### In vitro transcription

Reporter RNAs were generated using a previously established protocol (27, 28). Briefly, the linearized DNA was purified with the Wizard SV Gel PCR purification kit (Promega). For the synthesis of 5’-capped mRNA, reactions were carried out with the mMESSAGE mMACHINE T7 Transcription kit (Invitrogen). Capped-mRNAs were then polyadenylated with *E. coli* Poly-A polymerase (NEB) for 1 h at 37 °C. Synthesized mRNAs were purified by LiCl purification. The size and quality of the generating mRNAs were assessed by denaturing RNA gel electrophoresis.

### Human PURE in vitro translation

The human PURE translation was performed according to the established protocol (28, 47). Briefly, the human PURE cocktail (3.6 µL) was combined with 0.4 µL of reporter mRNAs (final concentration, 60 nM) and incubated at 32 °C for 6 h. In factor omission experiments, each translation factor was replaced with an equivalent volume of human PURE buffer (20 mM HEPES-KOH pH 7.5, 100 mM KCl, 10% (v/v) glycerol), maintaining the final concentration of each translation factor. Nluc assays were then performed as described previously (28).

### Cell culture

HEK293T cells were cultured in Dulbecco’s modified Eagle’s medium (DMEM; nacalai tesque) supplemented with L-glutamine/penicillin/streptomycin (2 mM L-glutamine, 100 U/mL penicillin, 0.1 mg/mL streptomycin; Sigma-Aldrich) and 10% (v/v) fetal bovine serum (ThermoFisher Scientific). Cells were incubated at 37 °C in 5% CO_2_. Cell density was maintained between 10% and 90%.

### Transfections and drug treatments

For in vivo C9-RAN luminescence assays, HEK293T cells were seeded in 96-well plates and transfected 24 h later at ~ 70% confluency. Transfections were performed using FuGene HD (Promega) at a 3:1 ratio to DNA, with 20 μL Opti-MEM containing 50 ng C9-RAN Nluc reporter DNA and 50 ng pcDNA-Fluc reporter per well. After 24 h, the medium was removed, and 100 μL of Glo Lysis Buffer was added. The plates were shaken for 1 min at room temperature. 30 μL of lysate was transferred to a new 96-well white plate. Luminescence measurements were taken using a Varioskan LUX Multimode Microplate Reader (ThermoFisher Scientific) and the Nano-Glo® Dual-Luciferase® Reporter Assay System (Promega). For knockdown experiments in 96 well plates, HEK293T cells were seeded at 30-40% confluency, and 2 pmol of siRNA per well was reverse-transfected using Lipofectamine RNAiMAX reagent (Invitrogen). After 24 h, cells were transfected with DNA reporters, and luciferase activity was measured as described above. siRNA sequences are listed in Supplementary Table S3. For overexpression experiments, Nluc reporter plasmids (20 ng/well) were co-transfected with Fluc plasmid (20 ng/well), and either eIF1A or eIF5B plasmid (200 ng/well) at a 1:10 ratio. After 24 h, luciferase activity was measured as described above. For C9-RAN reporter luminescence assays under eIF1A or eIF5B knockdown and ISR conditions, HEK293T cells were seeded and knocked down as described above for 24 h, followed by transfection with DNA reporters. Eighteen hours later, cells were treated with drugs for 6 h to induce ISR, followed by luciferase measurement as above. Drugs used: DMSO (Wako) and thapsigargin (ThermoFisher; final concentration, 2 μM).

For C9-RAN western blots, HEK293T cells were seeded in 12-well plates and transfected 24 h later at ~70% confluency. Transfections were performed using FuGene HD at a 3:1 ratio to DNA with 50 μL Opti-MEM containing 500 ng C9-RAN Nluc reporter DNA per well. Cells were lysed in 100 μL RIPA buffer supplemented with protease inhibitor (Roche) and PhosSTOP (Roche) 24 h post-transfection for 30 min at 4 °C. Cell lysates were then centrifuged at 15,000 × g for 15 min at 4 °C to remove debris, mixed with 4× SDS-sample buffer, and stored at −20 °C. For Western blots under eIF1A and eIF5B knockdown conditions, HEK293T cells were seeded at 30-40% confluency, and 20 pmol of siRNA per well was reverse-transfected using Lipofectamine RNAiMAX reagent (Invitrogen). After 24 h, cells (~80% confluency) were lysed as described above. For Western analysis following ISR activation, HEK293T cells were seeded at ~50% confluency. Twenty-four hours later (~80% confluency), cells were treated with drugs for 6 h to induce ISR, then lysed as described above. Protein concentrations for all Western blot samples were determined using Pierce™ BCA Protein Assay Kits (ThermoFisher).

### HEK293T cell lysate in vitro translation

HEK293T cell extracts were prepared using previously established protocols (49, 50). Briefly, HEK293T cells were washed with PBS and collected by trypsin treatment. The cell slurry was centrifuged, and the pellets were washed again with PBS. Cell pellets were then resuspended in an equal volume of A-buffer (10 mM HEPES-KOH (pH 7.5), 10 mM potassium acetate, 0.5 mM magnesium acetate, 5 mM dithiothreitol (DTT), and cOmplete protease inhibitor (Roche)). The suspension was incubated on ice for 45 min in a cold room, disrupted using a 26G syringe, further incubated for 10 min on ice, and centrifuged at 14,000 × g for 1 min at 4 °C to remove debris. The supernatant was collected and stored at −80 °C. For preparation of eIF1A KD lysate, HEK293T cells were seeded at 30-40% confluency in a 10 cm dish, and 200 pmol of siRNA were reverse-transfected using Lipofectamine RNAiMAX reagent (Invitrogen). After 24 h, cells were transferred to a 15 cm dish, and lysate was prepared as described above. For in vitro translation, HEK293T lysate (2.4 μL) was supplemented with Mix 1 (0.45 μL; 12 mM ATP, 8.2 mM GTP, 8.2 mM UTP, 8.2 mM CTP, 196 mM creatine phosphate, 1.2 mg/mL creatine kinase, 0.3 mM 20 amino acid mixture, 5 mM spermidine) and Mix 2 (1.65 μL; 97 mM HEPES-KOH pH 7.5, 290 mM CH_3_COOK, 8.6 mM MgOAc, 12.7 mM DTT). Reporter mRNAs were added at a final concentration of 3 nM, and the mixture was incubated at 32 °C for 1 h.

### Western Blotting

Protein samples were separated on an 11% WIDE range SDS-polyacrylamide system (nacalai tesque) for ~70 min and transferred to PVDF membranes. Membranes were blocked with 3% (w/v) skim milk in TBS-T, followed by incubation with primary antibodies listed in Supplementary Tables S4. After washing with TBS-T, membranes were incubated with HRP-conjugated anti-mouse IgG (Sigma-Aldrich) or anti-rabbit IgG (Sigma-Aldrich). Antibody reactions were performed using Can Get Signal (TOYOBO) when indicated. Chemiluminescence signals were detected using an LAS4000 imaging system (FujiFilm).

### Polysome profiling

HEK293T cells were seeded at 40-50% confluency in a 15-cm dish, and reverse transfection was performed using siRNA at a final concentration of 20 nM with Lipofectamine RNAiMAX reagent (Invitrogen). After ~24 h, cells were treated with 100 μg/mL cycloheximide (CHX, Wako) at 37 °C for 3 min to halt elongation, washed with PBS containing CHX, and collected using a cell scraper. Cells were lysed in low-salt lysis buffer (20 mM HEPES-KOH pH 7.5, 50 mM KCl, 10 mM MgCl_2_, 1% (v/v) Triton X-100, 1 mM DTT, 0.5% (w/v) sodium deoxycholate, 100 µg/mL cycloheximide, cOmplete protease inhibitor (Roche), PhosSTOP (Roche), and RNase Inhibitor Murine (NEB)) and centrifuged at 2,000 × *g* for 5 min. The supernatants were transferred to new tubes and centrifuged again at 13,000 × *g* for 5 min to remove fine debris. The resulting supernatants were layered onto 10-50% sucrose gradients prepared with low-salt buffer (same as the lysis buffer but excluding Triton X-100 and sodium deoxycholate) in Open Top Polyclear™ centrifuge tubes (14 × 89 mm, SETON) and centrifuged at 39,000 rpm for 2.5 h at 4 °C (Beckman OptimaL-90K, SW41-Ti). Fractions were collected using a Gradient Station (BIOCOMP) with a MICRO COLLECTOR AC-5700 (ATTO). Ribosome distribution was monitored by A_254_ absorbance using a BIO MINI UV MONITOR (AC-5200S, ATTO).

### Recombinant eIF1A-SE mutant protein expression and purification

Recombinant GST-tagged eIF1A-SE mutant protein was expressed in *E. coli* Rosetta 2(DE3) (Sigma # 71397-4) cultured in 2×YT medium supplemented with 20 μg/mL chloramphenicol and 100 μg/mL ampicillin. Protein expression was induced with 1 mM isopropyl-β-D-thiogalactopyranoside (IPTG) when the culture reached an optical density at 600 nm of 0.4-0.6. Cells were grown at 37 °C for 4-6 h, washed with PBS (nacalai tesque), and the pellets were stored at −80 °C. The frozen cell pellets were resuspended in lysis buffer (20 mM HEPES-KOH pH 7.5, 500 mM KCl, 2 mM DTT, 0.1%(v/v) Triton-X-100, 10%(v/v) glycerol, and cOmplete EDTA-free protease inhibitor cocktail tablet (Roche)), and lysed by sonication. Cell lysates were cleared by centrifugation for 30 min at 20,000 × *g*. The supernatant was incubated with Glutathione Sepharose 4B (Cytiva) for 1 h at 4 °C. The resin was washed with the same buffer and a high-salt wash buffer (20 mM HEPES-KOH pH 7.5, 1 M KCl, 2 mM DTT, 0.1%(v/v) Triton-X-100, 10%(v/v) glycerol). The buffer was then exchanged with PreScission cleavage buffer (50 mM Tris-HCl pH 7.5, 500 mM NaCl, 10%(v/v) glycerol, 0.1%(v/v) Triton-X-100, 1 mM EDTA, 1 mM DTT), and the GST-tag was cleaved on-column using PreScission protease (Cytiva) for 16 h at 4 °C. The flow-through was collected and applied to a PD-10 column (GE Healthcare) pre-equilibrated with human PURE buffer. After elution with 3.5 mL of the same buffer, the eluate was concentrated and stored at −80 °C.

### Quantification and statistical analysis

All statistical analyses were performed using custom R codes. Quantitative data are presented as mean ± standard deviation (S.D.) from at least three independent experiments.

## RESULTS

### Reconstituted translation system to screen potential regulators of RAN translation

Previously, we recapitulated C9-RAN translation using the human PURE system containing minimal translation factors, including eIFs, eEFs, and eRFs, and in vitro-transcribed mRNAs (Fig. 1A) (28). Using this system, we examined the role of specific translation factors in RAN translation by generating customized human PURE systems lacking individual factors and compared the results to canonical AUG codon translation (Fig. 1B, Supplementary Fig. S1). Omission of several translation factors inhibited translation, similar to effects observed in canonical AUG-initiated translation (Fig. 1C). Notably, depletion of eIF4F components—eIF4A (RNA helicase), eIF4B (eIF4A accessory protein), and eIF4G (scaffold protein)—led to a marked reduction in translation, suggesting that RAN translation, like canonical translation, proceeds via a cap-dependent mechanism, as previously reported (23–25, 27, 28). Conversely, removal of certain factors such as eIF1A, eIF5, and eIF5B significantly enhanced RAN translation (Fig. 1C). Previous studies have reported eIF5 as a promoter of RAN translation (35). However, our current findings suggest that eIF5 may function as a repressor, as its depletion enhanced RAN translation. We propose that this repressive effect is linked to eIF5’s role in facilitating eIF5B binding to the translation initiation complex under in vitro conditions (51). Thus, the absence of eIF5 may mimic the effect of eIF5B depletion. Based on these observations and the known roles of specific translation initiation factors, we propose that eIF1A and eIF5B act as regulatory factors in RAN translation, and further analyses were conducted to validate this hypothesis.

### eIF1A suppresses non-AUG codon initiation in RAN translation

Our translation factor screening using the human PURE system identified eIF1A as a candidate regulator. Previous studies have shown that eIF1A is involved in scanning for the optimal AUG codon (52–54). Based on this, we hypothesized that eIF1A represses non-AUG codon initiation in C9-RAN. To test this, we investigated whether eIF1A functions as a regulator of C9-RAN. First, we prepared HEK293T cell lysates with eIF1A knockdown (KD) (Supplementary Fig. S2A) and examined their response to C9-RAN. As observed in the human PURE system, translation from the GA(0) frame of C9-RAN was selectively enhanced in eIF1A-KD cells (Fig. 2A). Next, recombinant eIF1A was purified from *E. coli* and added to the cell lysates to assess its effect on non-AUG-initiated translation and RAN translation. Translation from a CUG codon was significantly repressed in a control reporter lacking repeats, where the AUG codon was replaced with a CUG codon (Fig. 2B). Similarly, translation from the GA(0) frame of C9-RAN was repressed by recombinant eIF1A, whereas this repression was abolished in a reporter in which the putative CUG start codon was mutated to AUG (Fig. 2B). In addition, eIF1A suppressed NOP56-RAN translation (AW(+1)) driven by a GGCCUG-71 repeat expansion (Fig. 2B). These findings suggest that eIF1A acts as a repressor of RAN translation by modulating initiation at non-AUG codons. To extend these observations, we knocked down or overexpressed eIF1A in HEK293T cells using a series of cell-based reporters (Supplementary Fig. S3). Upon eIF1A KD, translation from canonical AUG codons remained unchanged, whereas C9-RAN translation was enhanced across all frames (Fig. 2C, Supplementary Fig. S4A). Polysome profiling of eIF1A-KD cells revealed a slight increase in monosomes (Supplementary Fig. S4B, C), suggesting a global reduction in translation.

**Figure 2.**
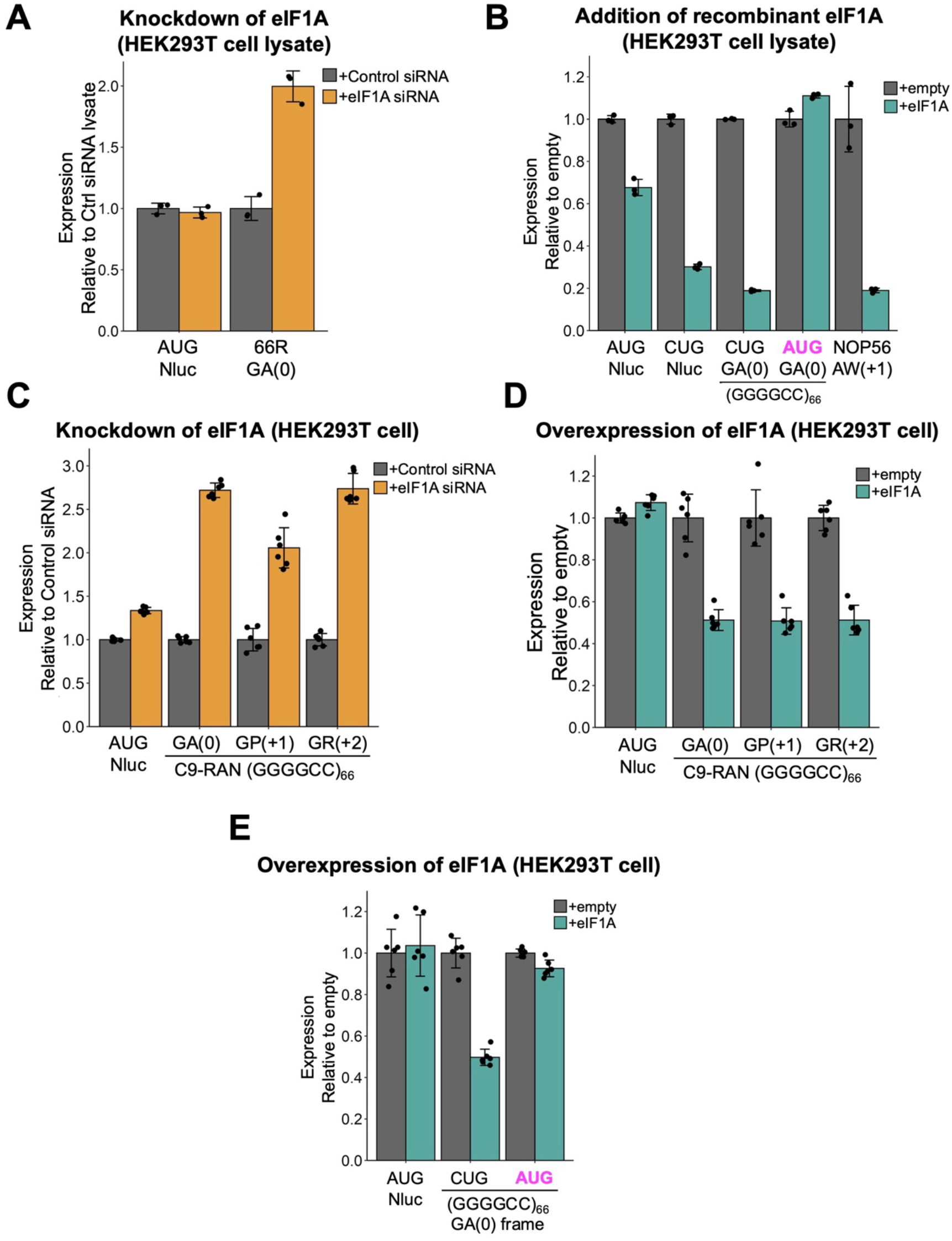
eIF1A modulates C9-RAN in a non-AUG codon-dependent manner. (**A**) Relative expression of indicated Nluc mRNA reporters in eIF1A knockdown (KD) HEK293T cell lysates, normalized to non-targeting control KD lysates (see also Supplementary Fig. S2). (**B**) Relative expression of Nluc reporters in HEK293T cell lysates supplemented with recombinant eIF1A (final concentration: 3 μM), normalized to lysates without eIF1A (vehicle control). (**C**) Relative expression of C9-RAN reporters in HEK293T cells 24 h after transfection with either eIF1A-targeting or non-targeting siRNAs. Nluc levels were normalized to non-targeting siRNA controls. (**D**) Relative expression of the indicated reporters in HEK293T cells co-transfected with C9-RAN reporters and an eIF1A expression plasmid. Nluc levels were normalized to the empty vector. (**E**) Relative expression of mutant C9-intron reporters in HEK293T cells co-transfected with mutant C9-RAN reporters and an eIF1A expression plasmid. Nluc levels were normalized to the empty vector. All graphs show mean ± SD from at least three to six independent experiments.

ISR is a well-established mechanism of translation inhibition in cells (Supplementary Fig. S5A) (55, 56). Since ISR is known to enhance RAN translation (24, 27, 30), we examined eIF2α phosphorylation to assess whether eIF1A KD activates ISR. However, eIF2α phosphorylation levels were not increased (Supplementary Fig. S5B), suggesting that RAN translation enhancement is not a secondary effect of ISR activation following eIF1A KD.

Next, eIF1A overexpression had no effect on translation from canonical AUG codons but repressed all frames of C9-RAN (Fig. 2D-E, Supplementary Fig. S6A). Notably, longer repeat lengths were also suppressed (Supplementary Fig. S6B, C). These effects were abolished when the CUG start codon in the GA(0) frame was mutated to a canonical AUG codon (Fig. 2E), indicating that eIF1A-mediated regulation of RAN translation depends on non-AUG initiation, consistent with the results from the human PURE and cell lysate systems.

To further investigate how eIF1A suppresses C9-RAN translation, we employed a bicistronic reporter system designed to assess IRES-like non-canonical initiation of C9-RAN (Supplementary Fig. S7A). The first cistron encodes Fluc via cap-dependent translation, followed by stop codons in all three frames. Downstream, either a GGGGCC repeat or HCV-IRES was inserted upstream of Nluc. In this system, Nluc is translated only through a cap-independent mechanism. eIF1A overexpression did not affect Nluc expression driven by HCV-IRES but significantly suppressed Nluc translation when preceded by the C9-RAN sequence (Supplementary Fig. S7B). These results indicate that eIF1A specifically inhibits C9-RAN translation without broadly affecting cap-independent initiation.

### Scanning function of eIF1A is associated with C9-RAN repression

Subsequently, we examined which specific regions of eIF1A are critical for repressing C9-RAN. eIF1A consists of an unstructured N-terminal tail (NTT), C-terminal tail (CTT), and an oligonucleotide-binding (OB) fold domain (Supplementary Fig. S8A). The CTT includes a scanning enhancer (SE) element that promotes the P_out_ conformation of the 43S scanning complex (52, 54). Mutations in this SE element have been shown to promote translation initiation from near-cognate codons in yeast (52, 54). We designed an SE mutant in which the predicted SE element in human eIF1A was substituted with alanine and purified the mutant protein. The SE mutant failed to suppress C9-RAN, and translation initiated from the putative CUG start codon in lysate-based translation (Fig. 3A). Similarly, overexpression of the SE mutant in HEK293T cells, where it was expressed at ~4.0-fold higher levels than endogenous eIF1A (Supplementary Fig. S8B), did not suppress C9-RAN in any reading frame, although a modest effect was observed in the GR (+2) frame (Fig. 3B).

**Figure 3.**
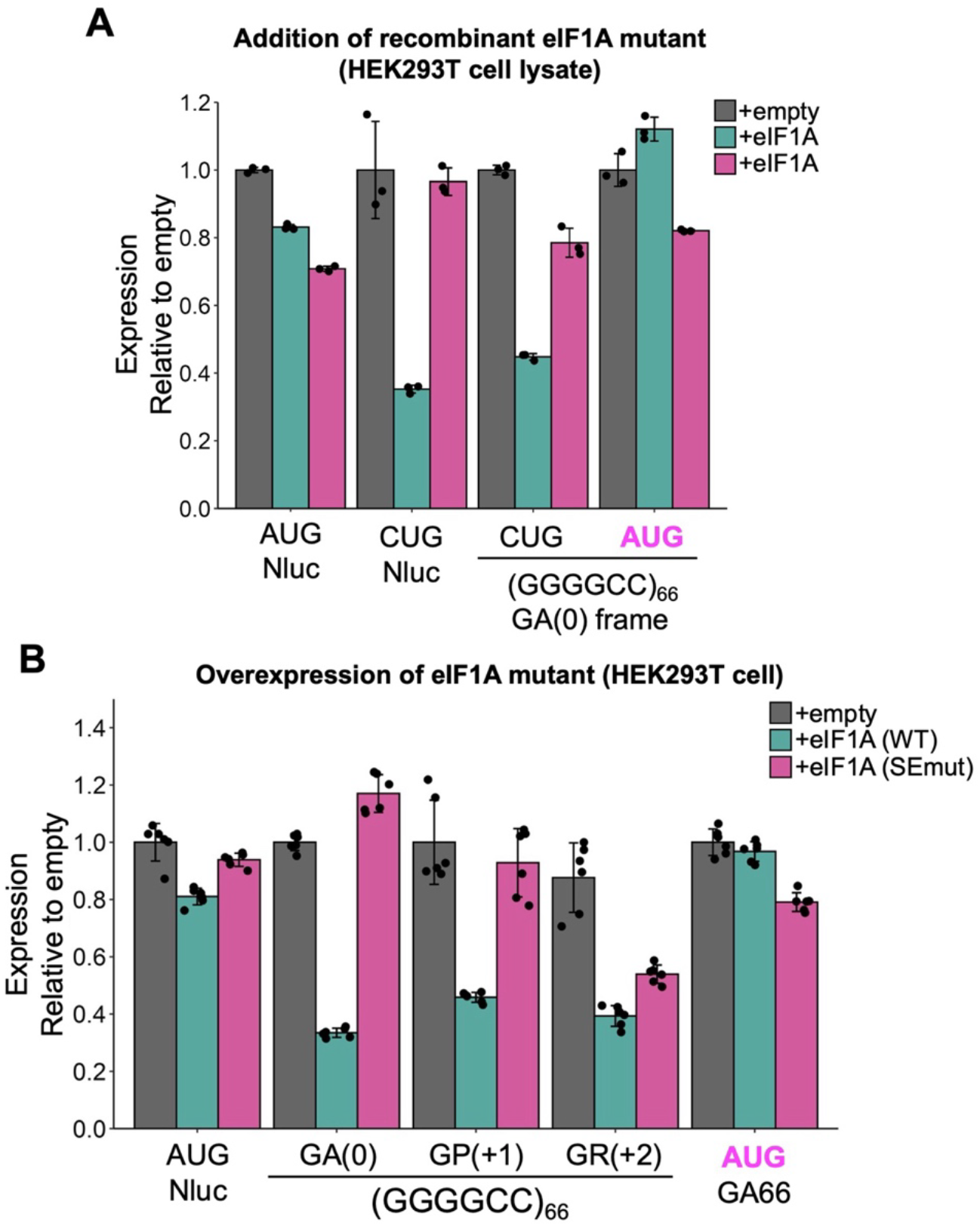
The C-terminal scanning enhancer (SE) region of eIF1A is essential for C9-RAN. (**A**) Relative expression of the indicated reporter in HEK293T cell lysates supplemented with wild-type or SE mutant eIF1A proteins (final concentration: 1 μM). Reporter mRNA was added to the lysates with each recombinant protein, and Nluc expression was normalized to the vehicle control. (**B**) Relative expression of the C9-RAN reporter in HEK293T cells co-transfected with eIF1A expression plasmids encoding either wild-type or SE mutant proteins. Nluc levels were normalized to the empty vector. All graphs show mean ± SD from at least three to six independent experiments.

The CTT of eIF1A also contains a contact site for eIF5B (residues 140-144; DIDDI) (57, 58), while the NTT mutation R13P disrupts the P_in_ state stability (59). We tested whether overexpression of these eIF1A mutants affects C9-RAN. The R13P mutant showed no significant impact (Supplementary Fig. S8C), suggesting that P_in_ state stability has minimal influence on C9-RAN. In contrast, the DIDDI mutant showed suppression of C9-RAN. Although the SE mutant weakly suppressed the GR(+2) frame (Fig. 3B), the DIDDI mutant suppressed all frames to a similar extent, suggesting that it affects C9-RAN through a mechanism distinct from that of the SE domain.

### eIF5B suppresses the initiation phase of C9-RAN via a mechanism distinct from eIF1A

Mutations at the eIF5B interaction site of eIF1A attenuated the C9-RAN suppression (Supplementary Fig. S8C) (58, 60). Based on this finding and our screening results using the human PURE system (Fig. 1C), we assumed that eIF5B might also function as a regulatory factor in C9-RAN. To test this, we examined eIF5B knockdown and overexpression in HEK293T cells, following the same approach as with eIF1A. Knockdown of eIF5B increased translation across all C9-RAN frames (Fig. 4A, Supplementary Fig. S9A), whereas overexpression suppressed C9-RAN in all frames (Fig. 4B, Supplementary Fig. S9B). Furthermore, when the GA (0) frame start codon was mutated to AUG, eIF5B-mediated suppression was abolished (Fig. 4C), indicating that, like eIF1A, eIF5B specifically inhibits non-AUG initiation.

**Figure 4.**
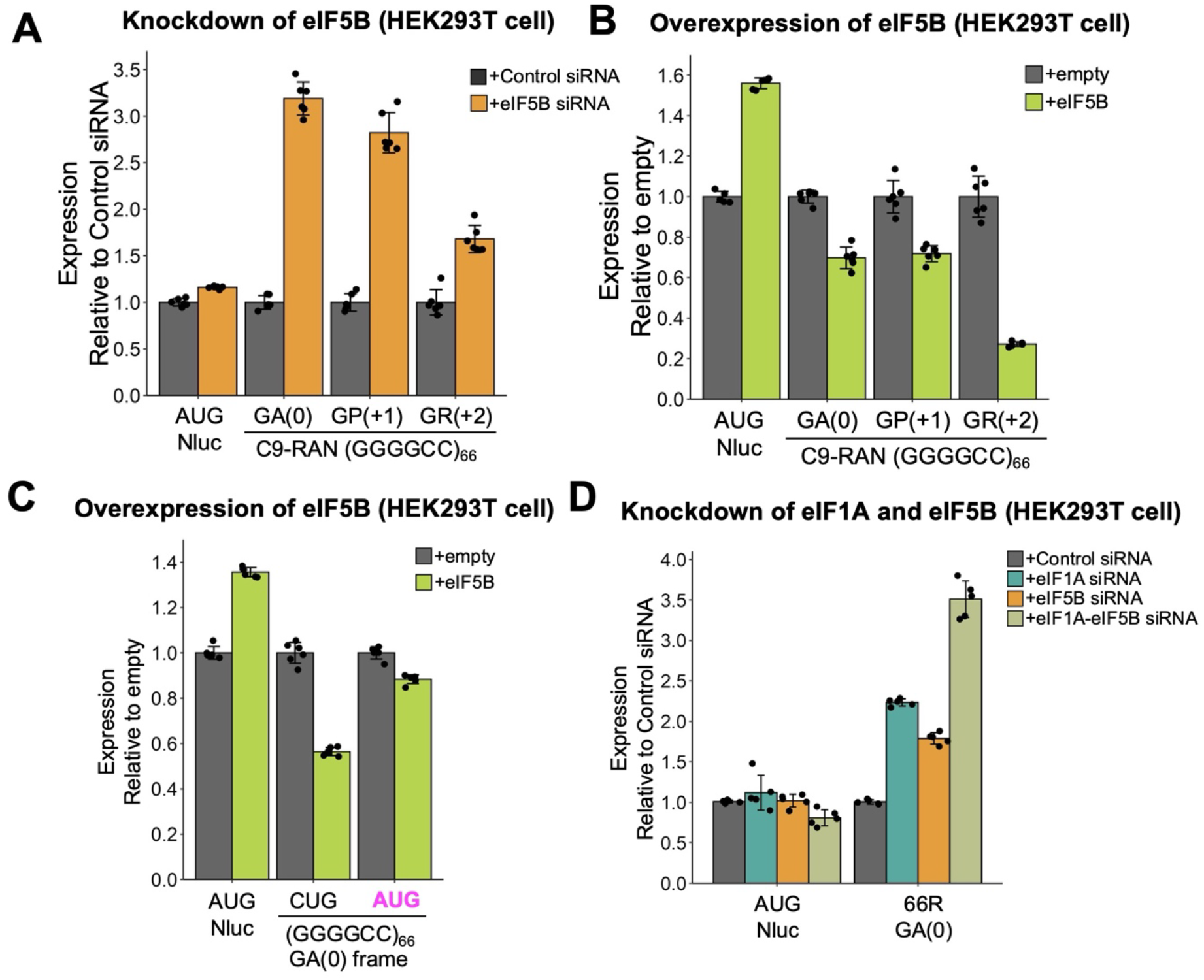
eIF5B acts as a C9-RAN repressor independent of eIF1A. **(A)** Relative expression of C9-RAN reporters in HEK293T cells, 24 h after transfection with either non-targeting or eIF5B-targeting siRNAs. Nluc levels were normalized to the non-targeting siRNA controls. **(B)** Relative expression of the indicated reporters in HEK293T cells co-transfected with C9-RAN reporters and the eIF5B expression plasmid. Nluc levels were normalized to the empty vector. (**C**) Relative expression of mutant C9-intron reporters in HEK293T cells co-transfected with the mutant C9-RAN reporters and the eIF5B expression plasmid. Nluc levels were normalized to the empty vector. (**D**) Relative expression of C9-RAN reporters in HEK293T cells, 24 h after transfection with non-targeting siRNA, individual knockdown siRNAs targeting eIF1A or eIF5B, or a combination of both (double knockdown). Nluc levels were normalized to the non-targeting siRNA controls (71). All graphs show mean ± SD from at least six independent experiments.

Given that eIF1A functions during scanning and eIF5B acts post-scanning (20– 22, 57), we postulated that they repress C9-RAN translation via distinct mechanisms. To test whether their effects are additive, we performed double knockdown of eIF1A and eIF5B in HEK293T cells. C9-RAN was further enhanced by the double knockdown compared to either single knockdown (Fig. 4D), supporting the idea that eIF1A and eIF5B independently regulate C9-RAN initiation.

### eIF1A contributes to the promotion of C9-RAN under ISR

As shown above, eIF1A knockdown in HEK293T cells did not activate ISR (Supplementary Fig. S5). However, ISR has been reported to enhance C9-RAN (24, 27, 30), though the mechanism underlying this enhancement remains unclear. To investigate whether eIF1A and/or eIF5B contribute to ISR-mediated promotion of C9-RAN, we treated eIF1A- or eIF5B-knockdown cells with thapsigargin (TG), a compound that induces ISR through ER stress (Supplementary Fig. S5A). In control siRNA-treated cells, TG-induced ISR increased C9-RAN across all frames, consistent with previous reports (Fig. 5). In contrast, TG-induced promotion of C9-RAN was abolished in eIF1A-KD cells, indicating that eIF1A is indispensable for ISR-dependent upregulation of C9-RAN. Conversely, TG still promoted C9-RAN in the eIF5B-KD cells. Since eIF1A is involved in scanning and eIF5B in post-scanning steps such as 60S subunit joining, these results suggest that proper scanning is critical for ISR-induced promotion of C9-RAN.

**Figure 5.**
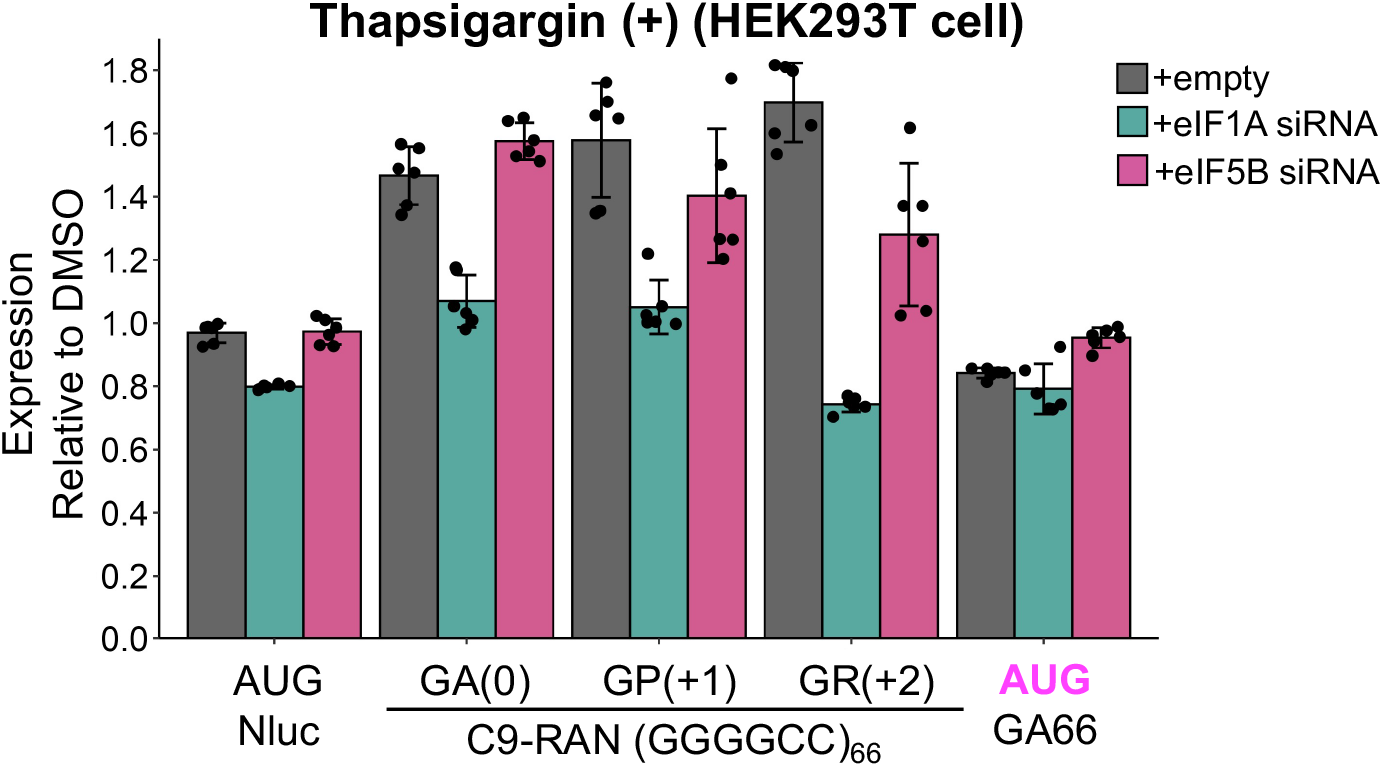
eIF1A is required for ISR-induced upregulation of C9-RAN. HEK293T cells were transfected with non-targeting, eIF1A, or eIF5B siRNAs for 24 h, followed by transfection with C9-RAN reporter plasmids. At 19 h after reporter transfection, cells were treated with 2 μM thapsigargin or vehicle (DMSO) for 5 h. Nluc expression was measured and normalized to vehicle-treated controls. Data represent mean ± SD from at least six independent experiments.

## DISCUSSION

In this study, we conducted a bottom-up screening for regulatory factors of C9-RAN translation using the human PURE system. Through this approach, we identified eIF1A and eIF5B as repressors of C9-RAN. We further validated their roles in lysate-based and cellular models, confirming that their regulatory effects are consistent across multiple experimental systems. This underscores the robustness of our bottom-up approach for identifying RAN translation regulators. eIF1A suppresses C9-RAN by regulating scanning to restrict initiation at non-AUG codons through its C-terminal SE region, while eIF5B, which is required for 60S subunit joining (post-scanning), would modulate C9-RAN. Both factors act at distinct stages of translation initiation to repress C9-RAN. Furthermore, our analysis in HEK293T cells under ISR conditions suggests that eIF1A is required for ISR-induced upregulation of C9-RAN. These findings reveal a functional interplay between eIF1A and eIF5B in controlling C9-RAN translation and suggest that proper scanning and ribosome assembly are essential for its regulation (Fig. 6).

**Figure 6.**
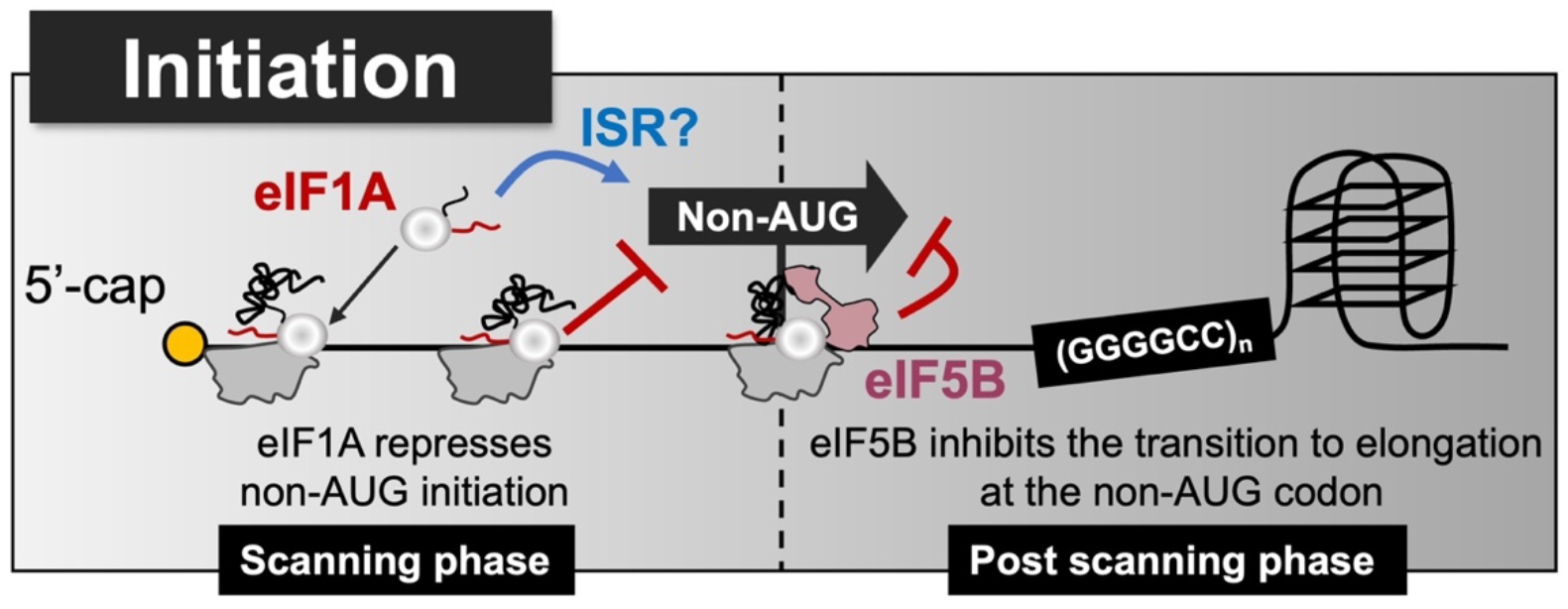
Plausible model of eIF1A- and eIF5B-mediated repression of C9-RAN. Details are described in the text.

eIF1A is essential for promoting scanning during translation initiation and is recognized as a key factor in ensuring accurate start codon selection (22, 52–54, 59). It is primarily associated with the 43S pre-initiation complex (PIC), where it regulates the transition between the P_in_ state (in which the PIC remains engaged with the mRNA) and the P_out_ state (in which scanning proceeds efficiently) through its N-terminal and C-terminal tails (NTT/CTT) (52, 54, 59). The SE region has been shown to play a critical role in maintaining the P_out_ state of the 43S PIC, ensuring that the scanning PIC accurately identifies the optimal start codon (52, 54). In yeast, mutations in the SE region have been reported to increase initiation at near-cognate start codons (52). Based on our findings, we propose that eIF1A suppresses C9-RAN by maintaining the fidelity of start codon selection. While eIF1 is also thought to contribute to the maintenance of the P_out_ state (61), our reconstituted system showed that eIF1 omission reduced both C9-RAN and canonical AUG-initiated translation, unlike eIF1A omission, which selectively enhanced C9-RAN. This suggests that eIF1 acts as a global regulator of scanning-dependent translation, regardless of start codon identity. However, it is noteworthy that eIF1 selectively suppresses FMR1 CGG-RAN translation (34), suggesting a potential role in regulating specific RAN translation events.

eIF1A has been identified in Drosophila as a factor that mitigates toxicity caused by arginine-rich DPRs, such as poly(GR) and poly(PR), which are translated via C9-RAN (46). However, our analysis using eIF1A-KD cells showed that poly(GR) production was not reduced, suggesting that eIF1A alleviates toxicity through a mechanism other than direct C9-RAN suppression. Previous studies in yeast have demonstrated that eIF1A plays a key role in maintaining start codon fidelity (52–54, 59). Therefore, differences in the relative abundance of translation factors and the systems used in various model organisms may account for the inconsistent effects observed with translation factor overexpression or knockdown. The failure to identify eIF1A as a C9-RAN regulator in previous studies is likely due to these system-specific differences. Conventional approaches using cell lysates or in vivo models may introduce indirect effects or obscure the direct regulatory role of eIF1A. In contrast, our screening with the human PURE system enabled precise control of translation factors, allowing us to identify eIF1A as a key regulator of C9-RAN.

eIF5B plays a critical role in the post-scanning phase of translation initiation by facilitating 60S ribosomal subunit joining, ensuring proper 80S ribosome assembly, and promoting the transition from initiation to elongation (20–22, 58, 60, 62). We found that, like eIF1A, eIF5B suppresses non-AUG initiation of C9-RAN. Unlike eIF1A, eIF5B is not predicted to have a major role in scanning (20–22). Given that C9-RAN initiates from near-cognate start codons, we propose that eIF5B acts as a checkpoint regulating 80S ribosome assembly at non-canonical initiation codons, thereby enforcing stricter translational control. Recent structural studies have shown that eIF1A and eIF5B cooperate to correctly position Met-tRNA_i_ in the P-site, reinforcing the fidelity of translation initiation (20–22). Indeed, eIF5B has been reported to enhance non-AUG translation in rabbit reticulocyte lysate when present in excess (63). eIF5B also interacts with eIF2A, which is involved in non-AUG initiation and in recruiting Met-tRNA_i_ to the ribosome (64). These findings suggest that eIF5B and eIF2A may cooperate to promote non-AUG translation. However, in our previous study, we did not observe the predicted cooperative effect of eIF5B and eIF2A (28). This discrepancy suggests that eIF5B-mediated regulation of non-canonical initiation may be context-dependent and influenced by additional cellular factors.

Both eIF1A and eIF5B act as suppressors of C9-RAN under normal conditions, but under ISR conditions, only eIF1A remains effective (32, 35, 38, 41). ISR generally suppresses global translation via phosphorylation of eIF2α while promoting translation of stress-responsive genes such as ATF4 (55, 56, 65). This upregulation occurs via leaky scanning, where upstream ORFs are bypassed, allowing translation of the main ATF4 ORF (66). A similar mechanism may operate in C9-RAN, where leaky scanning complexes accumulate upstream of the repeat sequence, increasing misrecognition of start codons and enhancing translation. In eIF1A-KD cells, these scanning complexes may fail to accumulate, preventing ISR-dependent upregulation of C9-RAN. This hypothesis could be further tested using translational complex profiling (TCP-seq) (67, 68). Alternatively, eIF1A KD may disrupt ISR-associated uORF-mediated translation dynamics, such as ATF4, resulting in the loss of ISR-responsive regulatory factors that selectively affect C9-RAN. In either case, our findings demonstrate that eIF1A is essential for ISR-mediated enhancement of C9-RAN.

A major strength of this study is the use of a bottom-up approach based on the reconstitution of RAN translation using the human PURE system. In cell lysates or cultured cells, knockdown or knockout of translation factors can cause secondary effects, making it difficult to assess their direct roles. In contrast, the human PURE system enables dissection of complex translation processes in a defined and minimal setting, eliminating secondary influences and providing a unique advantage for studying translation mechanisms. Using this reconstituted approach, we identified candidate factors essential for cell growth, which are typically difficult to detect in cell-based KD screening. While this minimal system does not reflect pathological conditions and is purely mechanistic, it complements conventional in vivo approaches. Engineered human PURE systems that modulate the balance of translation factors or mimic stress-induced translational environments will offer deeper insights into the molecular mechanisms of RAN translation.

## ACKNOWLEDGEMENTS

We thank the Center for Integrative Bioscience at Science Tokyo for DNA sequencing. We thank Dr. Petrucelli for kindly providing the pAG3-*C9orf72* intron1-(G_4_C_2_)_66_-3FT used in this study.

## FUNDING

This work was supported by MEXT Grants-in-Aid for Scientific Research (grant numbers JP26116002, JP18H03984, JP21H04763, and JP20H05925 to HT), AMED-

CREST under Grant Number JP21gm1410008 (to H.T.), JST, the establishment of university fellowships towards the creation of science technology innovation, JSPS KAKENHI (grant numbers JPMJFS2112 and 24KJ1067 to H. Ito), Uehara Memorial Foundation (to H.T.), Mitsubishi Foundation (to H.T.), and Daiichi Sankyo (to H.T.).

## AUTHOR CONTRIBUTIONS

H.Ito, K.M., Y.F., M.H. and S.S. performed experiments; H.Ito K.M. Y.F. Y.N. H.Imataka. and H.T. conceived the study, designed experiments, analyzed the results, approved the manuscript, and are accountable for all aspects of the work; H.T. supervised the entire project; H.Ito and H.T. wrote the manuscript with the help of other authors.

## CONFLICT OF INTEREST

The authors declare that they have no conflict of interest with the contents of this article.

## DATA AVAILABILITY

Data in this manuscript have been uploaded to the Mendeley Data repository (doi: 10.17632/vvdbs49mct.1).

## Notes

### Competing Interest Statement

The authors have declared no competing interest.

