## Supplemental figures for "Canonical translation factors eIF1A and eIF5B modulate the initiation step of repeat-associated non-AUG translation"

**Supplementary Figures S1-S9**

Supplementary Fig. S1: Schematic of bottom-up screening using a reconstituted human PURE system lacking individual canonical initiation factors.

Supplementary Fig. S2: Expression levels of eIF1A in eIF1A-KD cells.

Supplementary Fig. S3: C9-RAN reporters in HEK293T cells.

Supplementary Fig. S4: eIF1A KD reduces global translation level

Supplementary Fig. S5: eIF1A KD does not stimulate ISR.

Supplementary Fig. S6: eIF1A overexpression reduces C9-RAN in a repeat length-dependent manner in HEK293T cells.

Supplementary Fig. S7: eIF1A overexpression reduces C9-RAN levels in a bi-cistronic reporter.

Supplementary Fig. S8: The eIF5B contact site on eIF1A (DIDDI) is involved in C9-RAN repression.

Supplementary Fig. S9: Expression level of eIF5B in HEK293T cells

**Supplementary Tables S1-S4 (in separate Excel files)**

Supplementary Table S1: Plasmids list.

Supplementary Table S2: Oligonucleotides list.

Supplementary Table S3: siRNA sequences list.

Supplementary Table S4: Antibodies list.

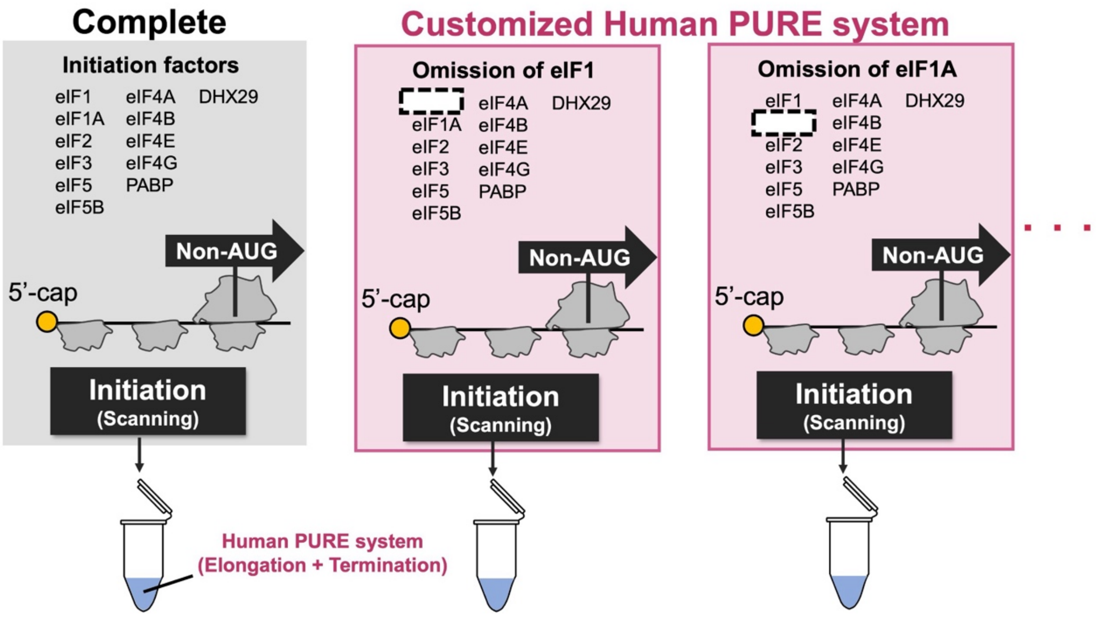


**Supplementary Fig. S1. Schematic of bottom-up screening using the human PURE system lacking individual canonical initiation factors.**

Schematic of a bottom-up screening approach using the human PURE system lacking individual canonical initiation factors.

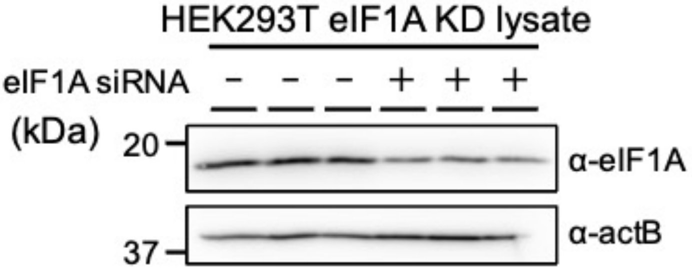


**Supplementary Fig. S2. Expression level of eIF1A.**

Western blotting analysis of lysates from non-targeting (-) and eIF1A siRNA (+) conditions, probed with an anti-eIF1A antibody to confirm reduced eIF1A expression. β-actin (actB) was used as a loading control.

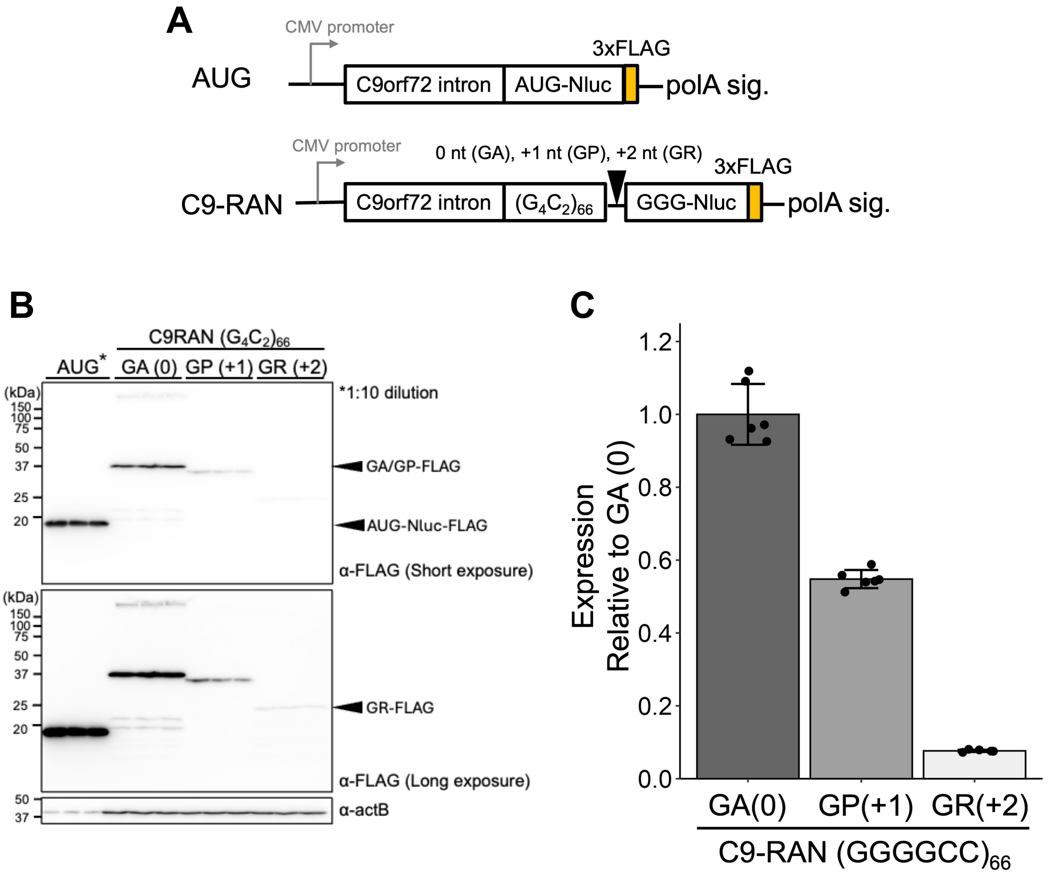


**Supplementary Fig. S3. C9-RAN reporter in HEK293T cells.**

*(A)* Schematic of C9-RAN Nanoluciferase (Nluc) reporters based on previously published constructs (35, 63–66). Nluc was used to measure translation efficiency.

*(B)* Anti-FLAG western blot of C9-RAN reporter expressed in HEK293T cells. The top and bottom panels show the same blot, with longer exposure in the bottom panel to detect the less abundant GR-FLAG. The AUG lane sample (*) was diluted 10-fold to match protein levels. β-actin (actB) was used as a loading control.

*(C)* Relative expression of C9-RAN reporters in HEK293T cells, normalized to GA(0). Data represent mean ± SD from at least six independent experiments.

A

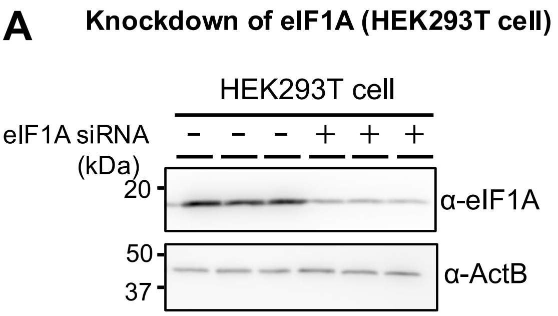


B

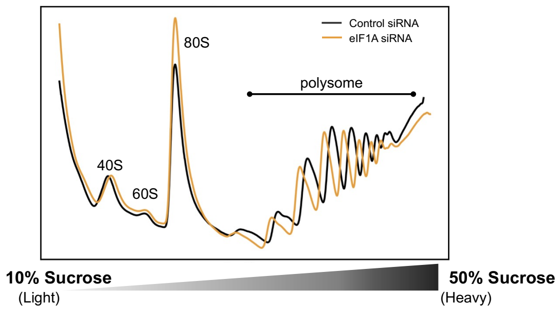


C

| Sample | Polysome/ Monosome ratio |
|---|---|
| Ctrl siRNA | 2.95 |
| eIF1A siRNA | 2.31 |

**Supplementary Fig. S4. eIF1A KD reduces global translation level.**

*(A)* Western blot showing the effect of eIF1A KD in HEK293T cells, 24 h after siRNA transfection and 24 h after reporter transfection. β-actin (actB) was used as a loading control.

*(B)* Polysome profiling of HEK293T lysates with non-targeting siRNA or eIF1A siRNA.

*(C)* Global translation activity (polysome/monosome ratio) calculated from the area under the plot in (B).

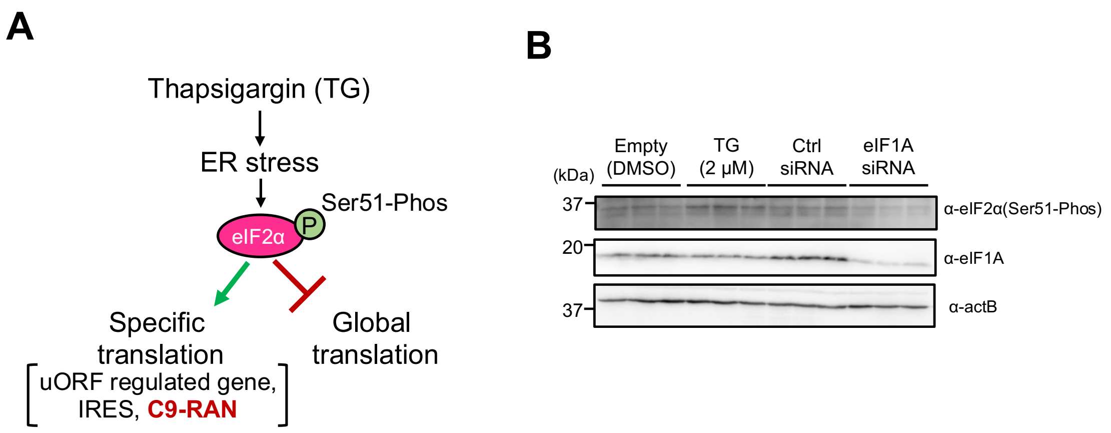


**Supplementary Fig. S5. eIF1A KD does not stimulate ISR.**

*(A)* Schematic of the integrated stress response (ISR) pathway (55, 56)

*(B)* Western blot showing the effect of eIF1A KD or 2 μM thapsigargin (Tg) treatment in HEK293T cells. β-actin (actB) was used as a loading control. n = 2.

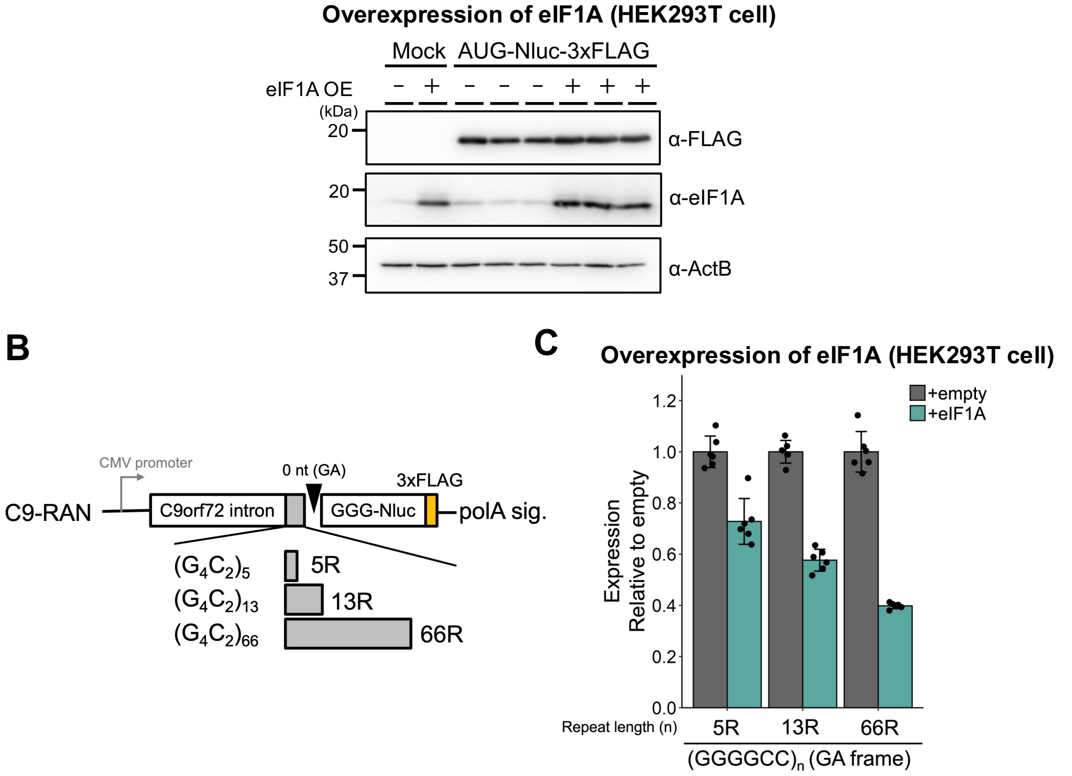


**Supplementary Fig. S6. eIF1A overexpression reduces C9-RAN in a repeat length-dependent manner in HEK293T cells.**

*(A)* Western blot showing the effect of eIF1A overexpression on AUG-Nluc-3xFLAG expression in HEK293T cells co-transfected with the C9-RAN reporter and eIF1A plasmid. Detection was performed using a FLAG tag. β-actin (actB) was used as a loading control.

*(B)* Schematic of the C9-RAN reporter containing different GA frame repeat lengths.

*(C)* Relative expression of C9-RAN with 5, 13, and 66 GA repeats in 293T cells co-transfected with empty or eIF1A expression plasmid. Nluc signals were normalized to the 5-repeat construct. Data represent mean ± SD (n = 6).

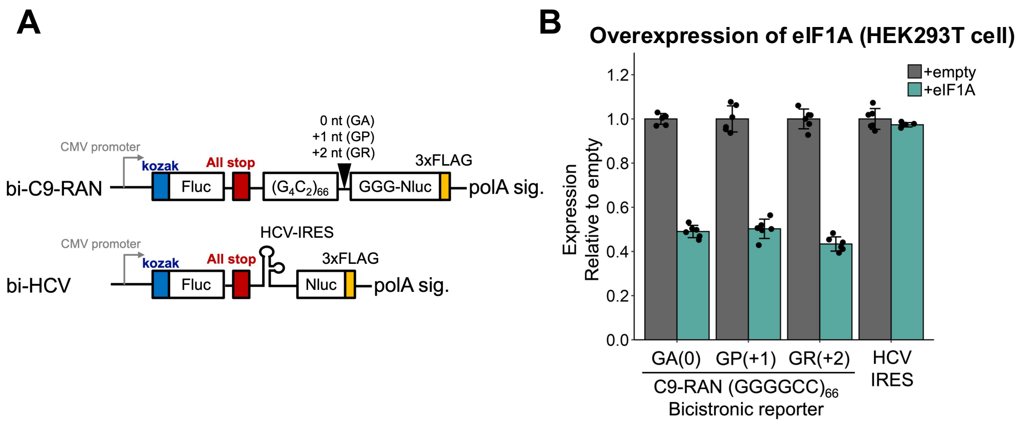


**Supplementary Fig. S7. eIF1A overexpression reduces C9-RAN levels in a bicistronic reporter.**

*(A)* Schematic of the bicistronic C9-RAN reporters for each reading frame.

*(B)* Relative expression of the indicated reporters in HEK293T cells co-transfected with bicistronic C9-RAN reporters and eIF1A expression plasmid. Nluc levels were normalized to the empty vector. Data represent mean ± SD (n = 6).

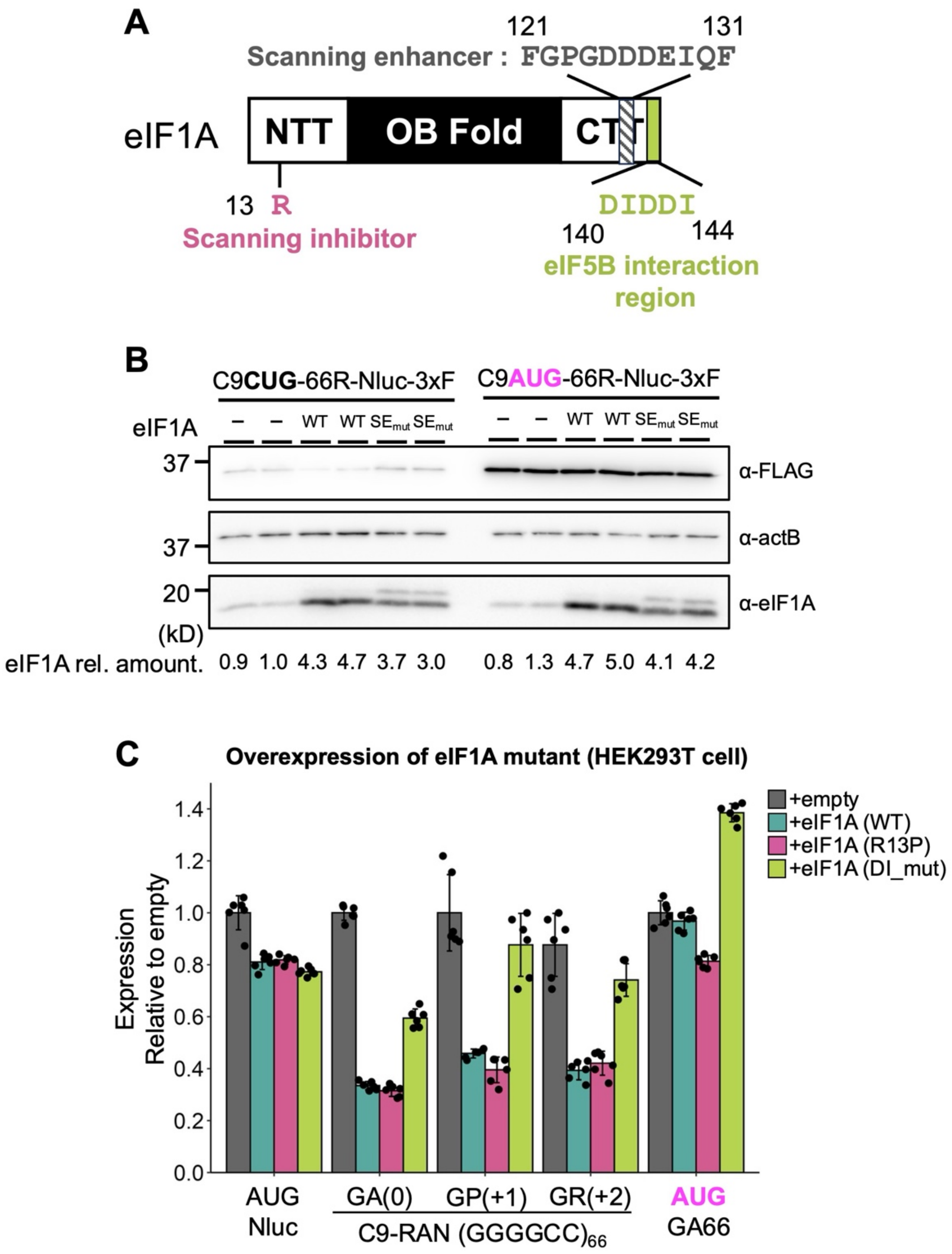


**Supplementary Fig. S8. The eIF5B contact site on eIF1A (DIDDI) is involved in C9-RAN repression.**

*(A)* Schematic representation of the domain structure of human eIF1A (52-54, 57-59).

*(B)* Western blot showing expression level of eIF1A from plasmids encoding wild-type or SE mutant proteins. C9-RAN detection was performed using a FLAG

antibody. eIF1A expression levels were normalized to the empty vector. β-actin (actB) was used as a loading control.

*(C)* Relative expression of the C9-RAN reporter in HEK293T cells co-transfected with eIF1A expression plasmids encoding wild-type, R13P, or DI mutant proteins. Nluc levels were normalized to the empty vector. Data represent mean ± SD (n = 6).

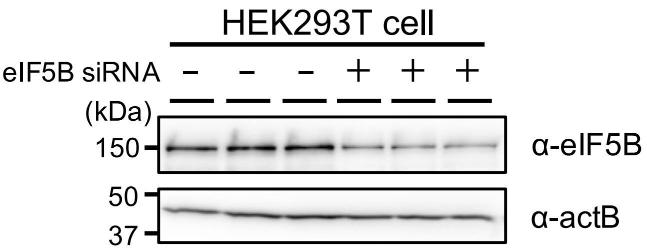


**Supplementary Fig. S9. Expression level of eIF5B in HEK293T cells.**
*(A)* Western blot showing the efficiency of eIF5B knockdown in HEK293T cells, 24 h after siRNA transfection. β-actin (actB) was used as a loading control.
